# Attention-related sampling of targets rhythmically alternates with increased susceptibility to co-occurring distractors

**DOI:** 10.1101/2025.08.07.669173

**Authors:** Zach V. Redding, Yun Ding, Ian C. Fiebelkorn

## Abstract

The Rhythmic Theory of Attention proposes that visual spatial attention is characterized by alternating states that promote either sampling at the present focus of attention or a higher likelihood of shifting attentional resources to another location. While theta-rhythmically (4–8 Hz) occurring windows of opportunity for shifting attentional resources might provide cognitive flexibility, these windows might also make us more susceptible to distractors. Here, we used EEG in humans to test how frequency-specific neural activity phasically influences behavioral performance and visual processing when high-contrast distractors co-occur with low-contrast targets. For trials with and without distractors, perceptual sensitivity at the cued target location depended on pre-stimulus theta phase (∼7 Hz) recorded at central electrodes. For trials with distractors, there was a greater increase in false alarm rates at the same theta phase associated with lower hit rates (i.e., during the proposed ‘shifting state’), confirming theta-rhythmically occurring windows of increased susceptibility to distractors. In addition to these phase-behavior effects at central electrodes, we observed phase-behavior effects at frontocentral and occipital electrodes that (i) only occurred on trials with distractors, (ii) peaked in the alpha-frequency range (∼9– 10 Hz) and (iii) were strongest at occipital electrodes that were contralateral to distractors. Alpha phase at these electrodes was also associated with fluctuations in the amplitude of distractor-evoked visual responses, consistent with an alpha-mediated gating of distractors. The present findings thus provide evidence for distinct theta- and alpha-mediated mechanisms of spatial attention that phasically modulate the influence of distractors on task performance.

**Significance:** The Rhythmic Theory of Attention proposes that spatial attention is characterized by alternating states that promote either sampling at the present focus of attention or a higher likelihood of shifting attentional resources to another location. These alternating attentional states are associated with dynamic changes in attention-related neural and behavioral effects, occurring on a timescale in the theta-frequency range (4–8 Hz). Although interdigitated windows for shifting attentional resources might provide critical cognitive flexibility, they might also lead to an increased susceptibility to distractors. Here, we demonstrate such rhythmic fluctuations in susceptibility to high-contrast distractors that co-occur with low-contrast visual targets. Rhythmic attention-related sampling—while perhaps preventing us from becoming overly focused on any single location—can lead to behavioral disadvantages.

## Introduction

The brain has limited processing resources. It therefore uses a collection of filtering mechanisms to enhance the processing of behaviorally important information and suppress the processing of distracting information. Spatial attention specifically refers to the filtering mechanisms through which the brain preferentially processes information at behaviorally important locations in space (Posner, 1980). The deployment of spatial attention is associated with changes in neural activity that improve behavioral performance (Carrasco, 2011; Moore and Zirnsak, 2017; Fiebelkorn and Kastner, 2019). These neural and behavioral effects, however, are not sustained during attentional deployment. Growing evidence instead indicates that attention-related effects fluctuate over time, on a theta-rhythmic timescale (i.e., at 4–8 Hz) (VanRullen et al., 2007; Busch and VanRullen, 2010; Landau and Fries, 2012; Fiebelkorn et al., 2013; Song et al., 2014; Dugué et al., 2015, 2016; Landau et al., 2015; Re et al., 2019). Fluctuations in attention-related effects have been linked to theta-rhythmic neural activity in the large-scale network that directs spatial attention and exploratory movements (i.e., the ‘Attention Network’) (Fiebelkorn et al., 2018, 2019; Helfrich et al., 2018; Spyropoulos et al., 2018; Gaillard et al., 2020).

The Rhythmic Theory of Attention proposes that the waxing and waning of attentional effects reflects alternating states associated with either sampling at the present focus of attention (i.e., a ‘sampling state’) or a higher likelihood of shifting attentional resources to another location (i.e., a ‘shifting state’) (Landau and Fries, 2012; Dugué et al., 2016; Fiebelkorn and Kastner, 2019; Senoussi et al., 2019; Benedetto et al., 2020). In support of this ‘shifting state’, exploratory behaviors, such as eye movement in primates, have similarly been linked to theta-rhythmic neural activity (Otero-Millan et al., 2008; Bosman et al., 2009; Tomassini et al., 2015, 2017; Hogendoorn, 2016; Lowet et al., 2016; Wutz et al., 2016). Frequently occurring windows of opportunity for shifting attentional resources could prevent us from becoming overly focused on any single location in space, thereby providing critical cognitive flexibility. On the other hand, rhythmically occurring shifting states could make us more susceptible to distracting information (i.e., behaviorally irrelevant information). Previous auditory research, for example, has demonstrated that the vulnerability of working memory to distractors fluctuates at low frequencies (Wöstmann et al., 2019; Lui et al., 2023). Here, we tested whether frequency-specific neural activity phasically influences susceptibility to distractors during attention-related sampling of the visual environment, specifically when spatially predictable, high-contrast distractors co-occur with spatially predictable, low-contrast targets.

In addition to hypothesized links between theta-band activity and distractor susceptibility, alpha-band activity (9–14 Hz) has been repeatedly associated with the suppression of sensory processing (Jensen and Mazaheri, 2010; Foxe and Snyder, 2011). Higher alpha power typically occurs within neural populations that are processing behaviorally irrelevant sensory information (Foxe et al., 1998; Worden et al., 2000; Thut, 2006) and is also associated with lower cortical excitability (Haegens et al., 2011; Spaak et al., 2012; Bonnefond and Jensen, 2015; Dougherty et al., 2017). For example, Haegens and colleagues (2011) demonstrated that higher alpha power was associated with lower spike rates. Despite these often-observed relationships between alpha-band activity and the suppression of sensory processing, research examining whether alpha-mediated mechanisms of spatial attention can be actively deployed to suppress spatially predictable distractors has provided mixed results (Foster and Awh, 2019; Bonnefond and Jensen, 2025). We therefore also tested whether pre-stimulus alpha-band activity—like pre-stimulus theta-band activity—modulates behavioral performance and visual processing when spatially predictable, high-contrast distractors co-occur with spatially predictable, low-contrast targets. The present findings provide evidence for distinct theta- and alpha-mediated mechanisms of spatial attention that both phasically modulate the influence of distractors on task performance. Moreover, these findings confirm a key prediction of the Rhythmic Theory of Attention: despite being behaviorally disadvantageous, there are theta-rhythmically occurring windows of increased susceptibility to distractors.

## Methods

### Subjects

Forty individuals with normal or corrected-to-normal vision and no history of neurological disorders participated in the experiment (26 females, 14 males; mean age, 24.1 years). All subjects provided informed consent. The study was conducted in accordance with protocols approved by the Research Subjects Review Board at the University of Rochester.

### Apparatus and stimuli

Subjects were seated in a comfortable chair in a sound- and light-attenuated chamber. Chair and table heights were adjusted so that the subject could comfortably use padded chin and forehead rests. A foot stool was provided if needed. Subjects were instructed to remain as relaxed as possible during trials to minimize noise in the EEG attributable to muscle activity. Task contingencies were controlled with custom Presentation (Neurobehavioral Systems) scripts run on a Dell Precision 5820 desktop computer with a 27-inch Acer Predator XB2 LCD monitor (1,920 × 1,080 pixels at 240 Hz) positioned 57 cm from the subject’s eyes. Figure 1 depicts the primary experimental display which consisted of a grey background with a light grey fixation square (0.5 × 0.5 dva) at the center of the screen and four light-grey placeholder circles of 4 dva centered at 6 dva from fixation and 45°, 135°, 225°, and 315° from vertical (i.e., one in each quadrant). Cues consisted of a solid border (outside diameter of 4.5 dva) around placeholder stimuli. Target cues were dark grey and distractor cues were orange or blue.

**Figure 1.**
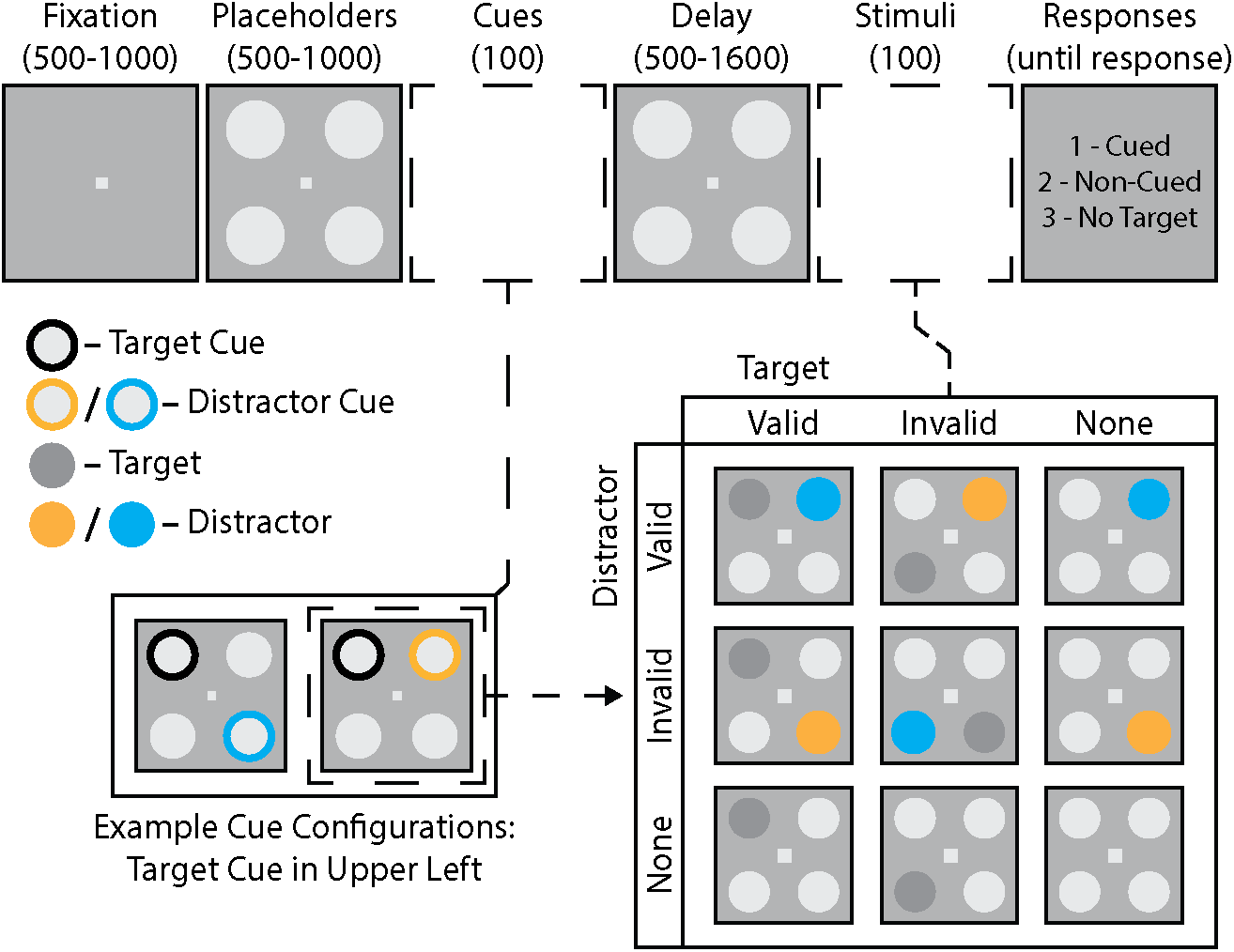
Experimental task. Trials began with variable fixation (500–1000 msec) and placeholder (500– 1000 msec) intervals, followed by two spatial cues (100 msec) on opposite sides of the visual field to indicate the likely locations of both a subsequent near-threshold target and a salient distractor (100 msec). Cue validity was 70% for both cue types. Targets and distractors were presented after a variable delay (500–1600 msec). Stimulus displays could include (i) both a target and a distractor, (ii) a target only, (iii) a distractor only, or (iv) neither a target nor a distractor. The number pad on a computer keyboard was used to indicate the presence of a target at the cued location, a target at a non-cued location, or no target.

Targets were near-threshold, grey, circular patches, 4 dva in diameter. Target luminance was adjusted throughout the task to maintain hit rates (HR) between 50% and 75% and to keep false alarm rates (FAR) below 30%. Distractors were highly salient, circular patches, 6 dva in diameter, and drawn in orange or blue (same colors as distractor cues).

### Behavioral task

Figure 1 illustrates the task. Participants pressed the left button on a computer mouse to begin each trial, causing a fixation square to appear for 500–1000 msec. Four placeholders then appeared for 500–1000 msec. Next, subjects were presented with a cue display consisting of a target cue (black) at one of the two positions on either side of the visual field and a distractor cue (randomly orange or blue) at one of the two positions on the opposite side. Cues remained on the screen for 100 msec. The subsequent display consisted of the four placeholder stimuli and lasted between 500–1600 msec, with the duration selected randomly from a uniform distribution. After the delay period, stimuli were presented for 100 msec. The stimulus configuration could consist of either i) four placeholder stimuli (i.e., catch trials; ∼17.6% of trials), ii) a target stimulus and three placeholder stimuli (∼23.5%), iii) a distractor stimulus and three placeholder stimuli (∼23.5%), or iv) a target, a distractor, and two placeholder stimuli (∼35.3%). 70% of all targets/distractors appeared at cued locations (i.e., valid), while 30% of targets/distractors appeared at non-cued locations (i.e., invalid). Distractor color (orange or blue) was randomly selected for each trial and independent of cue color. Following the offset of stimuli, placeholders returned to the screen for a brief period (300 msec). Participants were then shown a response screen with instructions to press 1 on the keyboard if they detected a target at the cued location, press 2 if they detected a target at a non-cued location, or press 3 if they did not detect a target. This display remained until participants responded.

Each subject completed at least 20 practice trials followed by six blocks of 170 trials for a total of 1,020 trials. Subjects were instructed to maintain fixation without blinking for the duration of each trial (i.e., the period beginning with the subject pressing the mouse button and ending with onset of the response screen). Fixation was monitored using an infrared eye-tracking camera (EyeLink 1000 Plus, SR Research). If fixation was broken (> 2 dva), the trial was aborted, and a warning message appeared, instructing subjects to begin the next trial. If trials were aborted in this way, additional trials were added to the block to ensure the total number of completed trials for each condition was consistent across subjects.

### EEG Recording and Pre-Processing

EEG data were collected using a BioSemi ActiveTwo system (BioSemi) with 128 active Ag/AgCl electrodes. Data were digitized at 2,048 Hz and downsampled to 512 Hz off-line. All electrode impedances were kept below 20 kΩ. To ensure consistent placement of the EEG cap, the vertex electrode (A1) was placed at 50% of the distance between the inion and the nasion and between the tragus on the left and the right ears. EEG data were analyzed using the FieldTrip toolbox (Oostenveld et al., 2011) and custom scripts written in MATLAB (R2023a). Off-line, continuous EEG data were re-referenced to the average of all 128 channels, high-pass filtered (fourth-order Butterworth with 0.1 Hz cutoff), and then segmented into epochs relative to stimulus onset (−2 to 0.5 s). Bad channels were visually identified during recording sessions. Artifacts related to blinks and eye movements were excluded from the data by design, as task performance was contingent on maintaining fixation (i.e., trials with a blink or saccade were aborted). Data were visually inspected and any trials exceeding a ± 100 µV threshold during analysis periods (e.g., delay period or ERP window) were interpolated using a distance-weighted average of the four nearest good electrodes, if the average distance was < 5 cm (otherwise data were excluded). Trials without all 128 electrodes were excluded from analyses and subjects with greater than 50% of these trials were excluded entirely, resulting in 8 subjects being excluded (n = 32).

### Behavioral Analysis

Based on visual inspection of response time histograms for individual subjects, trials with a response time less than 400 msec or greater than 2000 msec were excluded from these analyses. Paired samples t-tests were used to test for effects of target cueing on HR, FAR, and d-prime (D’). To isolate target cueing effects, only trials with no distractor were included. For calculations of FAR, trial counts were adjusted to account for the relative probability of targets at cued and non-cued locations (75% vs 25%). Paired samples t-tests were also used to test for effects of distractor presence and distractor cueing on HR, FAR, and D’. To isolate distractor and distractor cueing effects, only trials with valid targets were included in calculations of HR and only trials with responses at cued target locations were included in calculations of FAR. Cohen’s d was used to express effect sizes for all behavioral tests.

### ERP Analyses

ERP analyses were limited to the time window from −100 to 350 msec relative to distractor onset. This time window captures the sensory response to distractors without including activity evoked by response screen onset. Time series were baseline corrected using the period from −100 to 0 msec and averaged across trials. Only trials without targets were included in these analyses. Trials were randomly subsampled to match counts across conditions for each subject. Occipital channels A10/B7 were used for all ERP analyses, consistent with past work on visual responses to distractors (e.g.,

Hickey et al., 2009; Redding and Fiebelkorn, 2024). For analyses measuring the effects of pre-stimulus phase (see below) on distractor-evoked visual responses, we applied a Hilbert transform to the broadband filtered signal (1-55 Hz) and took the absolute value to capture non-phase-aligned, broad-band power. This procedure allowed us to baseline trials, which could not be done confidently for phase-aligned signals, as pre-stimulus binning based on phase creates systematic differences in the baseline of raw voltage signals. Cluster-based permutation tests were performed to compare distractor-evoked responses across conditions while controlling for multiple comparisons (Luck, 2014). Briefly, a t-test was performed to compare responses at each timepoint. We then calculated the sum of differences for clusters of consecutive timepoints at which a significant difference was observed (α = 0.05). Next, we shuffled the condition assignment across trials and repeated this procedure to identify the maximum sum of differences for a representative null cluster. This procedure was repeated 5000 times to form a null distribution against which we compared the sum of differences in observed clusters of significant timepoints. Observed clusters with a sum greater than 95% of null cluster sums were considered significant.

### Lateralized Phase-Behavior Analyses

Trials with a response time less than 400 msec or greater than 2000 msec were excluded from these analyses. For each condition (no distractor and valid distractor), trials were binned based on frequency-specific, pre-stimulus phase measurements. Phase was measured on each trial by taking the angle of the complex output produced by convolving the signal with a frequency-specific Morlet wavelet (3-55 Hz), with a varying number of cycles (2 cycles for 3-8 Hz, increasing logarithmically from 2 to 5 cycles for 9-55 Hz). Subsequent analyses focused on frequencies from 3-25 Hz. We only included trials with a delay period of at least 1000 msec to avoid capturing cue-evoked activity. Wavelets were fit for each frequency at a time point that positioned the last point of the wavelet immediately prior to stimulus onset, to avoid capturing stimulus-evoked activity. Performance measures were calculated for overlapping phase bins with a width of 120 degrees (e.g., 0–120 degrees) that were shifted forward in 10-degree steps (e.g., 10–130 degrees, 20–140 degrees, etc.). In contrast to prior work (Fiebelkorn et al., 2013, 2018; Abdalaziz et al., 2023), we leveraged the opposition of target and distractor cues to test for lateralized oscillations with functional links to target detection. In other words, we always measured phases with respect to the positions of relevant cues. This was achieved by flipping channels for trials with the distractor cue appearing on the right side, so that channels on the right side are always contralateral to the distractor cue (and ipsilateral to the target cue). This procedure was repeated to generate phase-behavior functions, spanning all phases, for each frequency-by-channel pair, which were then averaged across participants. Here, we predicted that the averaged phase-behavior functions would have a characteristic shape, with a peak separated from a trough by approximately 180 degrees (i.e., approximating a one-cycle sine wave) (see Figure 5). Following this prediction, a discrete Fourier transform was applied to the phase-behavior function for each frequency-by-channel pair, and the second component—representing a one-cycle sine wave—was kept. The amplitude of this component reflects both how closely the function approximated a one-cycle sine wave and the effect size, thereby providing a single value representing the strength of the phase-behavior relationship (Fiebelkorn et al., 2013, 2018; Abdalaziz et al., 2023). Phase-behavior analyses were applied to trials with (i) valid targets/responses at cued locations and (ii) either no distractor or a valid distractor. These analyses were not performed for invalid target/distractor conditions due to low trial counts.

### Topographic Analyses

To compare the topographies of phase-behavior effects, we adapted an approach used to test for differences between ERP topographies (Murray et al., 2008). For each condition, the strength of phase-behavior functions (i.e., one-cycle sine fit) was z-normalized across channels. We then calculated the square root of the mean of squared differences between the normalized values to obtain a single value representing the global dissimilarity between the topographies for the two conditions (Lehmann and Skrandies, 1980). Condition labels were then shuffled at the subject level and global dissimilarity was re-calculated. This process was repeated 5,000 times to create a reference distribution of global dissimilarity values. Pairs of topographies with global dissimilarity greater than 95% of all null values were considered significantly different.

### Analyses of Alpha Power

Only trials with delay periods longer than 900 msec were included in analyses of delay-period alpha power to avoid effects from cue-evoked visual responses. Alpha frequency can vary substantially between individuals (Haegens et al., 2014); therefore, peak alpha frequencies were determined for each subject. Identification of subject-specific alpha peak frequencies used data from channels A10/B7 which correspond to those used in prior work on the role of posterior alpha in attentional selection (e.g., Kelly et al., 2006; Wang et al., 2019; Redding and Fiebelkorn, 2024). Data were isolated from −500 to 0 msec relative to stimulus onset, and irregular resampling autospectral analysis (Wen and Liu, 2016) was used to obtain oscillatory power spectra (2–30 Hz) at electrodes contralateral to cued distractor locations. Data were padded with zeros to obtain a frequency resolution of 0.1 Hz. For each subject, local maxima in the alpha-band (8–15 Hz) were identified after averaging spectra across trials. Subject-specific alpha frequency was used for subsequent analyses of alpha power. For each trial, power was measured during the period from -500 to 0 msec relative to stimulus onset by applying a Hanning taper and discrete Fourier transform to the signal and taking the squared magnitude of the complex output. Data were padded with zeros to achieve 0.1 Hz resolution. Pre-stimulus alpha power was used to calculate a lateralization index using the following formula:

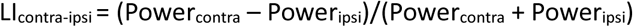

For trials with a distractor cue on the right side, lateralization indices for all channels were flipped across the midline before combining with data for trials with the distractor cue on the left side. Trials were binned by alpha lateralization (median split) at the occipital channel (A10/B7) contralateral to distractor cues and paired samples t-tests were used to test for an effect on task performance.

## Results

### Behavioral Effects

Here, our primary goal was to test whether oscillatory mechanisms of spatial attention influence the effects of a distractor on visual-target detection. To do this, we presented spatially predictable targets with spatially predictable distractors (Figure 1), using spatial cues with 70% validity. Prior to measuring the influence of oscillatory mechanisms on behavioral performance, we tested (i) whether the distractor interfered with visual-target detection (Wöstmann et al., 2022) and (ii) whether participants utilized the spatially informative target and distractor cues. Figure 2 shows that the presence of a distractor interfered with visual-target detection at cued target locations (relative to trials without a distractor), both reducing HR [*t*(31) = 4.312, *P* < .001, *d* = .762] and increasing FAR [*t*(31) = 3.035, *P* = .005, *d* = .537]. Combining HR and FAR, the presence of a distractor also reduced D’ [*t*(31) = 7.125, *P* < .001, *d* = 1.26], confirming that distractors were associated with reduced perceptual sensitivity at cued target locations.

**Figure 2.**
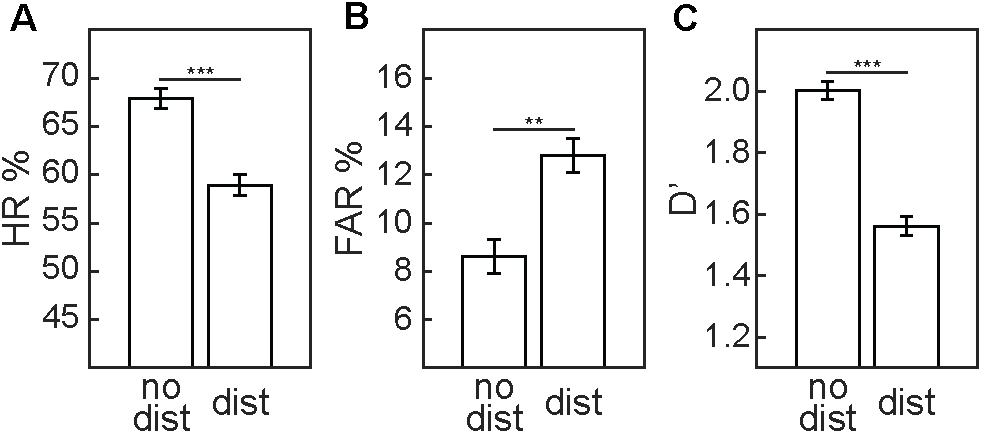
Distractors impaired task performance. ***A***, Mean HR for trials with no distractor and trials with a distractor. ***B***, As in ***A***, but for mean FAR. ***C***, As in ***A***, but for mean D’. All error bars indicate ±1 SEM. ** *P* < .01; *** *P* < .001.

We next tested whether spatially informative target and distractor cues were associated with better visual-target detection, relative to invalidly cued trials. Figure 3 illustrates both target- and distractor-cueing effects. Here, HR was significantly increased for validly cued targets, relative to invalidly cued targets [*t*(31) = 7.381, *P* < .001, *d* = 1.31], but there was no significant effect of target cueing on FAR [*t*(31) = 1.037, *P* = .308, *d* = 0.183]. Combining HR and FAR, D’ was significantly higher at cued target locations, relative to non-cued target locations [*t*(31) = 4.46, *P* = .001, *d* = 0.788]. Distractor cues were associated with a similar pattern of results. That is, HR was significantly increased when there was a validly cued distractor, relative to when there was an invalidly cued distractor [*t*(31) = 4.304, *P* < .001, *d* = .761], but there was no significant effect of distractor cueing on FAR [*t*(31) = 1.167, *P* = .252, *d* = .154].

**Figure 3.**
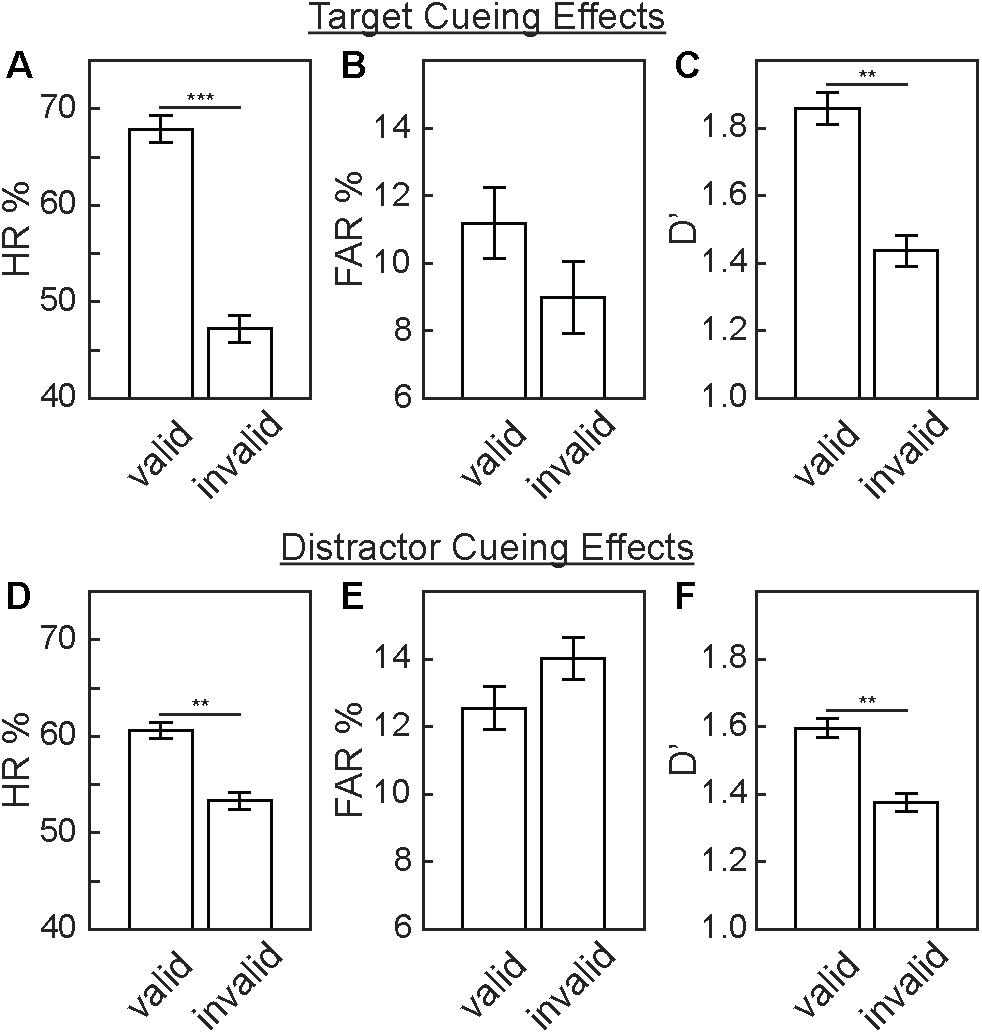
Spatially informative target cues and distractor cues improved task performance. ***A***, Mean HR for trials with valid targets and trials with invalid targets. ***B***, Mean FAR for responses at valid (i.e., cued) locations and invalid (non-cued) locations. ***C***, Mean D’ for valid/cued locations and invalid/non-cued locations. ***D***, Mean HR for trials with valid distractors and trials with invalid distractors. ***E***, As in ***D***, but for mean FAR. ***F***, As in ***D***, but for mean D’. All error bars indicate ±1 SEM. ** *P* < .01; *** *P* < .001.

Combining HR and FAR, D’ at the cued target location was significantly higher when there was a validly cued distractor, relative to when there was an invalidly cued distractor [*t*(31) = 4.065, *P* < .001, *d* = .719] (Figure 3D-F). These behavioral results confirm that participants utilized the spatially informative target and distractor cues to both enhance target processing at likely target locations and suppress distractor processing at likely distractor locations.

### ERP Evidence for Cue-Related Suppression of Distractor Processing

After confirming behavioral evidence of cue-related distractor suppression, we tested for neurophysiological evidence of such suppression. Figure 4A depicts grand-averaged ERP waveforms in response to either validly or invalidly cued distractors that were presented in the contralateral hemifield. There is a prominent P1 component, consistent with processing in extrastriate areas (Clark and Hillyard, 1996). Figure 4B depicts the difference wave associated with these distractor-evoked potentials (i.e., invalid minus valid). Cluster-based permutation tests comparing the visual responses to validly and invalidly cued distractors revealed a significant cluster from 136.5 to 152.1 msec after distractor onset (*P* = .016). The topography of the difference wave during this temporal window is consistent with a suppressive effect on sensory processing in occipital cortex (Figure 4C). Here, we flipped the topography from trials when the distractor occurred on the right side of the visual field, such that the right side of the present topography represents the visual response at electrodes that were contralateral to distractors (i.e., regardless of whether the distractor was presented to the right or to the left of central fixation) and the left side of the present topography represents the visual response at electrodes that were ipsilateral to distractors. These neurophysiological results provide further evidence that participants utilized the spatially informative distractor cue to suppress distractor processing. The low-contrast target did not elicit clear evoked responses, so we were unable to test for neurophysiological evidence of cue-related target enhancement.

**Figure 4.**
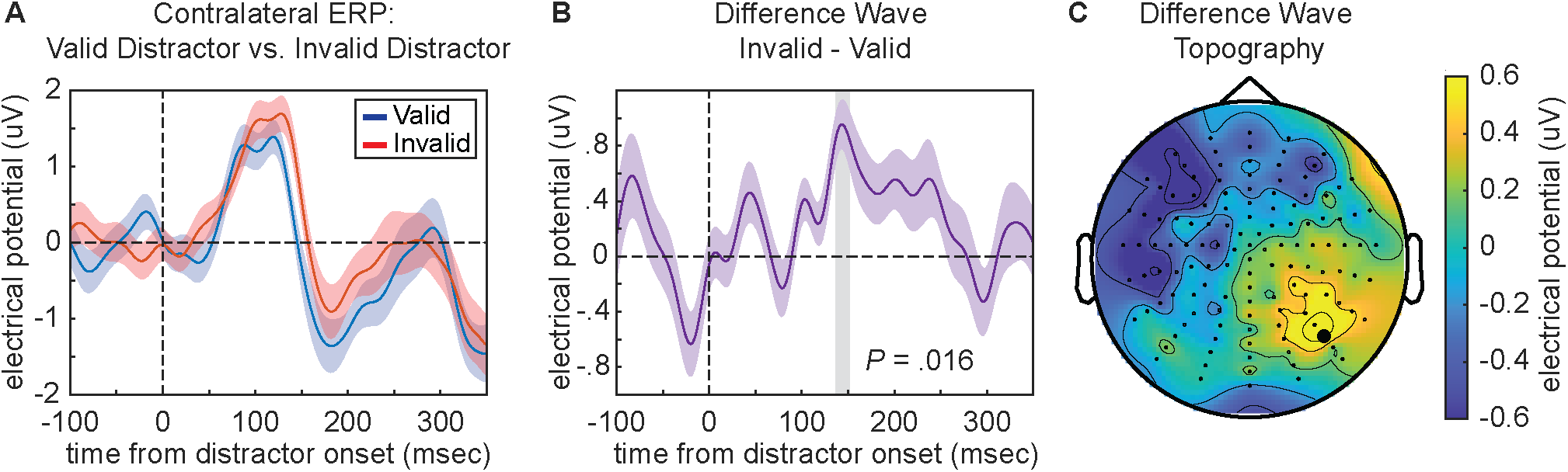
ERP evidence for suppression of distractors appearing at cued locations. ***A***, Grand average ERPs from occipital channels contralateral to valid distractors (blue), invalid distractors (red). ***B***, Grand average difference wave after subtracting valid distractor responses from invalid distractor responses. The time window of significant difference between conditions is indicated with a grey bar (136.5 to 152.1 msec). All error bars indicate ±1 SEM. ***C***, Topography of the difference wave averaged over the significant time window. The electrode used for ERP analyses is depicted with a bold dot.

### Distinct Oscillatory Mechanisms Phasically Modulate the Influence of Distractors on Task Performance

We next tested our primary research question, measuring the link between the pre-stimulus phase of frequency-specific neural activity and visual-target detection at the cued target location (Figure 5). We specifically measured these phase-behavior relationships with and without a validly cued distractor. Figure 6 displays significant phase-behavior relationships—based on cluster-based statistics (see Methods)—across behavioral measures and experimental conditions. For trials with and without a distractor, the phase of theta-band activity significantly modulated HR and D’ (Figure 6A, C). For trials without a distractor, these effects were strongest at ∼7 Hz, with topographies of the D’ effects revealing a central cluster and broad effects across bilateral frontal channels, with only frontal channels contralateral to the distractor cue reaching statistical significance (Figure 6D). FAR, in comparison, was only linked to theta phase on trials with a distractor (Figure 6B). Notably, the phase associated with the poorest HR was also associated with the highest distractor-related increase in FAR (Figure 7B). These results are consistent with the proposal that both visual target detection and distractor susceptibility fluctuate as a function of attention-related, theta-rhythmic sampling (Fiebelkorn and Kastner, 2019). We did not have enough trials to investigate phase-behavior relationships in the context of invalid targets and distractors.

**Figure 5.**
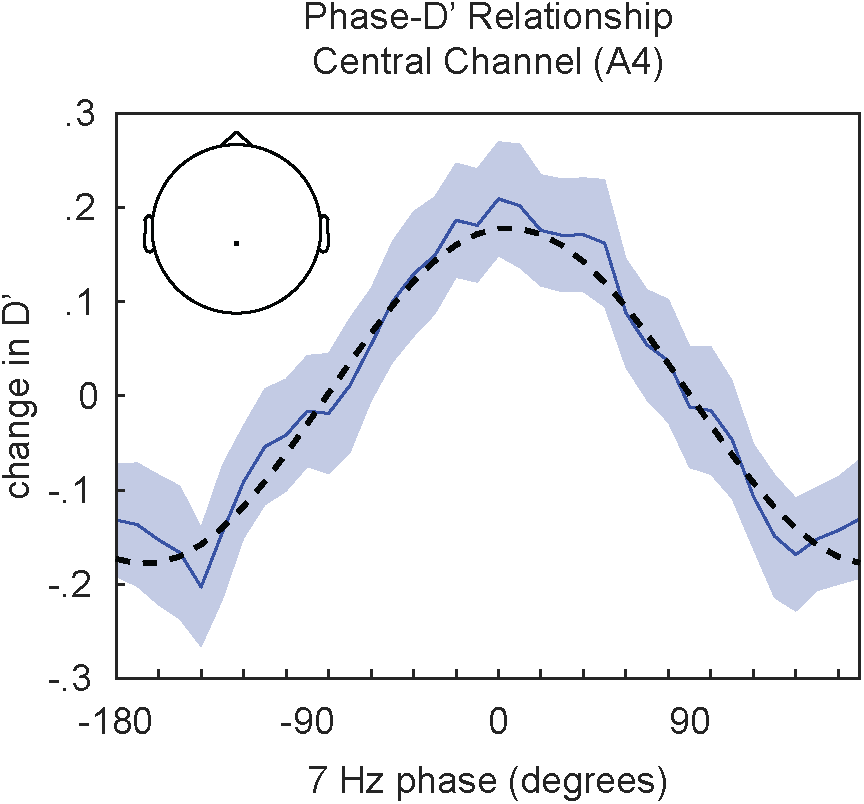
Example of phase-behavior analysis. Normalized behavioral measures are plotted as a function of the pre-stimulus phase of ongoing EEG oscillations. This example depicts the relationship between 7 Hz phase and D’ (solid blue line). Error bars indicate ±1 SEM. The amplitude of a fitted, one-cycle sine wave (dashed line) indicates the strength of the phase-behavior relationship. Inset depicts the electrode associated with this phase-behavior function.

**Figure 6.**
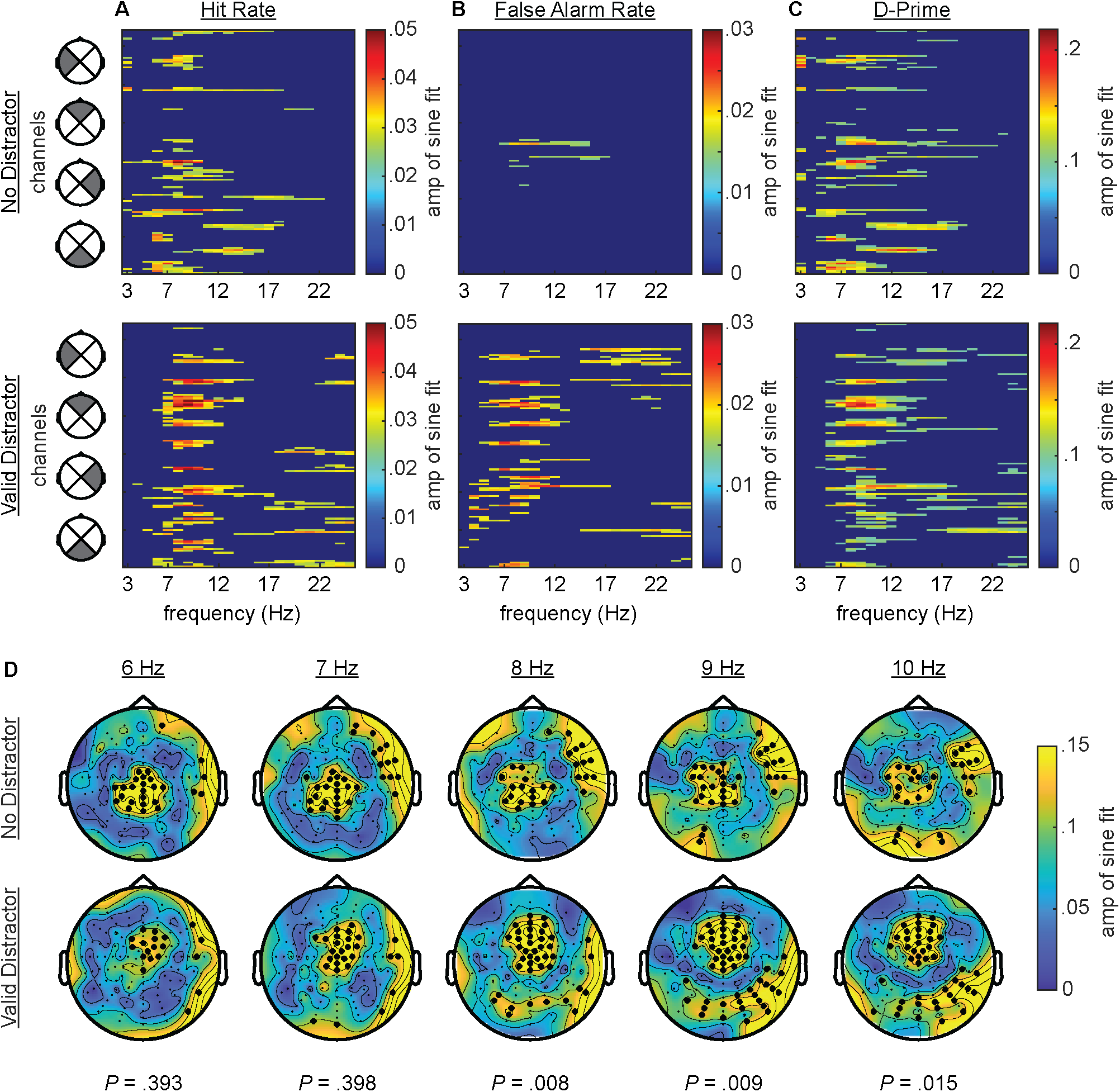
Phase-behavior relationships. ***A***, Plots depicting the strength of statistically significant (*P* < .05) phase-HR relationships as a function of frequency (3–25 Hz) across all 128 channels (statistically insignificant results were set to zero). The top row shows effects for trials with no distractor and the bottom row shows these effects for trials with a valid distractor. ***B***, As in ***A***, but for phase-FAR relationships. ***C***, As in ***A***, but for phase-D’ relationships. ***D***, Scalp topographies show the effect size for relationships between D’ and phase for frequencies from 6–10 Hz (columns) with significant channels indicated by bold dots. The top row shows effects for trials with no distractor and the bottom row shows effects for trials with a valid distractor. *P*-values for tests comparing these topographies across the no distractor and valid distractor conditions are listed below, with significant values (*P* < .05) indicating differences between the conditions.

**Figure 7.**
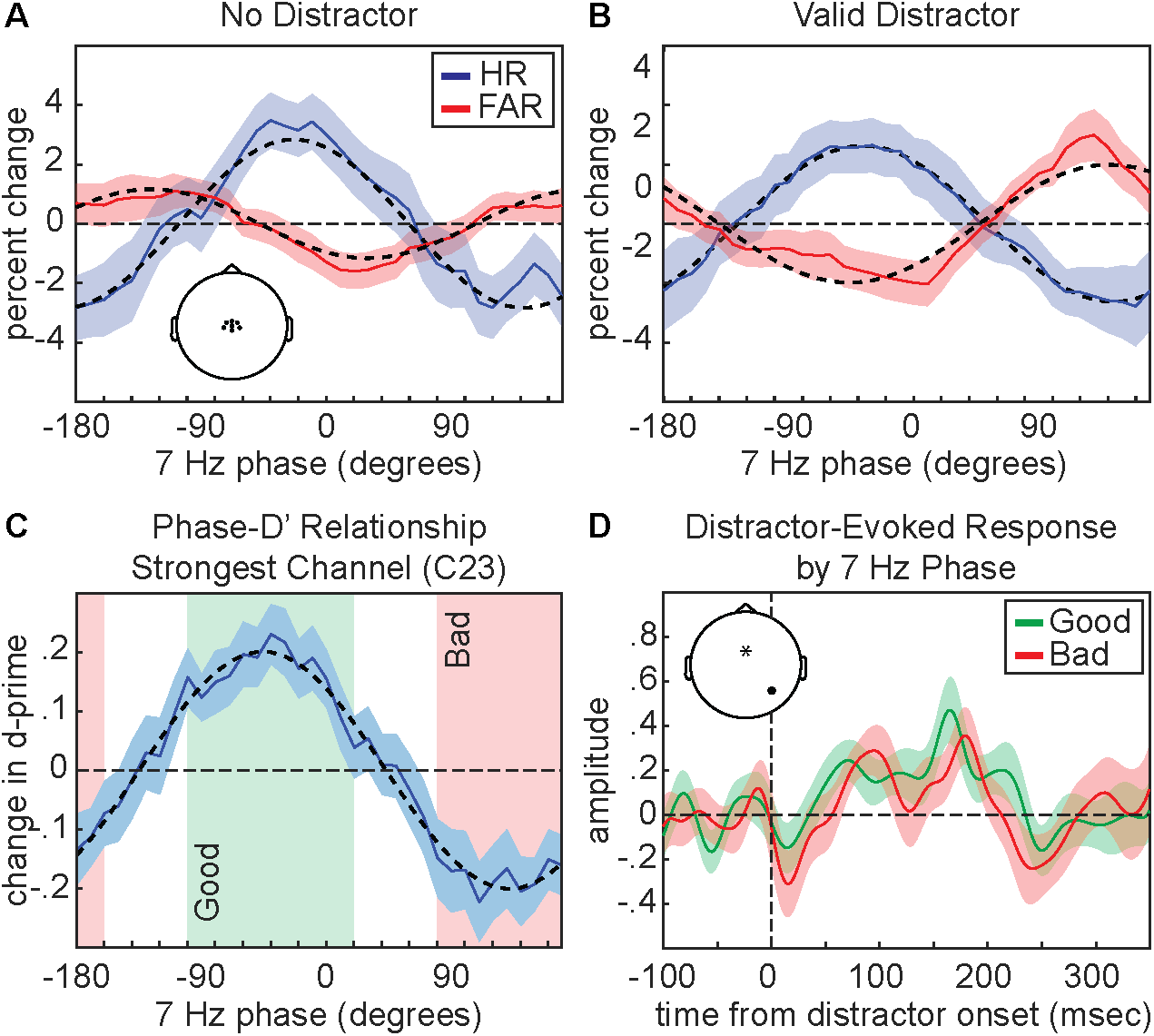
Task performance depends on pre-stimulus theta phase. ***A***, Phase-HR (solid blue line) and phase-FAR (solid red line) relationships for 7-Hz phase using trials with no distractor. These effects were averaged across central channels that were significant for both conditions (depicted in inset). ***B***, As in ***A***, but for trials with a valid distractor. ***C***, Phase-D’ relationship for the channel (C23) demonstrating the strongest effect on trials with a valid distractor (solid blue line). This function was used to determine ‘good’ (green box) and ‘bad’ (red box) phase bins. ***D***, Distractor-evoked broadband (1-55 Hz) response as a function of central 7-Hz phase (‘good’ vs. ‘bad’). Inset depicts the locations of the channel used for phase measurements (indicated by an asterisk) and the channel used for broadband responses (indicated with a dot). All error bars indicate ±1 SEM.

For trials with a distractor, all behavioral measures (i.e., HR, FAR, and D’) were also linked to the phase of higher frequency oscillations in the alpha band, with the strongest effects at ∼9-10 Hz (Figure 6A-C, Figure 8A-B, and Supplemental Figure 1). Topographies of the D’ effects revealed alpha-specific frontocentral and occipital clusters, with the occipital effects being stronger at electrodes that were contralateral to the distractor cue (Figure 6D). As with the ERP analyses, we flipped the topographies from trials when the distractor cue occurred on the right side of the visual field, such that the right side of the present topographies represent phase-behavior relationships at electrodes that were contralateral to distractor cues and the left side of the present topographies represent phase-behavior relationships at electrodes that were ipsilateral to distractors cues. The alpha phase associated with better behavioral performance at the frontocentral cluster (Figure 8B) is opposite from the alpha phase associated with better behavioral performance at the occipital cluster (Supplemental Figure 1). These antiphase results could reflect opposite ends of an alpha-related dipole, or perhaps a recent proposal that top-down distractor suppression is mediated by antiphase alpha-band activity in distinct sensory and downstream detection areas (Ursino, 2025).

**Figure 8.**
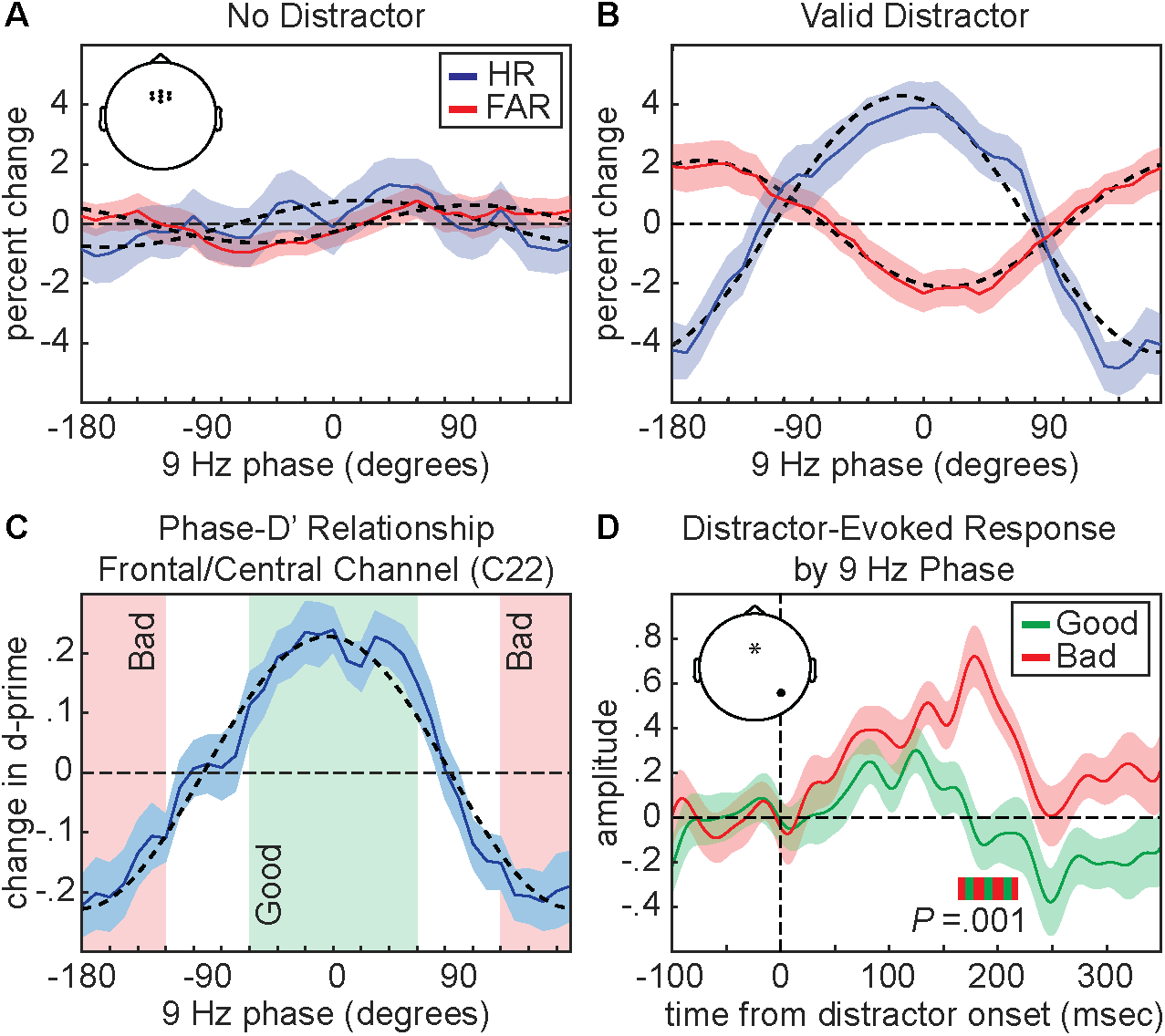
Distractor suppression depends on pre-stimulus alpha phase. ***A***, Phase-HR (solid blue line) and phase-FAR (solid red line) relationships for 9-Hz phase using trials with no distractor. These effects were averaged across frontocentral channels that were significant for trials with a valid distractor (depicted in inset). ***B***, Same as ***A***, but for trials with a valid distractor. ***C***, Phase-D’ relationship for the channel (C22) demonstrating the strongest effect on trials with a valid distractor (solid blue line). This function was used to determine ‘good’ (green box) and ‘bad’ (red box) phase bins. ***D***, Distractor-evoked broadband (1– 55 Hz) response as a function of frontocentral 9-Hz phase (‘good’ vs. ‘bad’). Inset depicts the locations of the channel used for phase measurements (indicated by an asterisk) and channel used for broadband responses (indicated with a dot). All error bars indicate ±1 SEM. Horizontal green and red bar indicates the time window for the significant difference between conditions (163.8 to 218.5 msec).

To further test how phase-behavior relationships differed between trials with and without distractors, we statistically compared topographies across the conditions (for frequencies from 6–10 Hz). These tests (no distractor vs. valid distractor) revealed no differences at 6 Hz (*P* = .393) and 7 Hz (*P* = .398) but significant differences at 8 Hz (*P* = .008), 9 Hz (*P* = .009), and 10 Hz (*P* = .015). The present results are therefore consistent with theta-dependent effects that occurred on trials with and without distractors and alpha-dependent effects that only occurred on trials with distractors (Figure 6D).

Finally, we tested whether pre-stimulus theta and alpha phase were associated with fluctuations in distractor-evoked visual responses. Here, we used broad-band power (from 1–55 Hz), rather than grand-averaged ERPs (see Methods). Using the channel with the strongest relationship between pre-stimulus phase and D’ on trials with a distractor (i.e., channel C23 for 7 Hz and channel C22 for 9 Hz), we binned trials by phase (+/- 60 degrees of the midpoint of bins with the best and worst behavioral performance) to create ‘good’ and ‘bad’ phase conditions (Figures 7C and 8C). While there was no effect of theta phase (Figure 7D) on distractor-evoked responses, there was a significant effect of alpha phase on distractor-evoked responses (Figure 8D). That is, the response to valid distractors was weaker when presented during the ‘good’ alpha phase relative to the ‘bad’ alpha phase, with a cluster-based permutation approach (see Methods) revealing a significant cluster from 163.8 to 218.5 msec after distractor onset (*P* = .001). These results are consistent with an alpha-mediated gating of distractor processing (Jensen and Mazaheri, 2010).

### Occipital Alpha Power Does Not Influence Suppression of Distractors at Expected Locations

While alpha-band activity has been repeatedly linked to the suppression of sensory processing, evidence for a relationship between alpha-band activity and distractor suppression has been mixed (Bonnefond and Jensen, 2025). In the previous analysis, the results demonstrated a significant relationship between pre-distractor alpha phase and subsequent behavioral and neurophysiological responses (Figures 6 and 8). Here, we tested whether pre-stimulus alpha power was also linked to behavioral performance on trials with distractors. We specifically tested whether alpha-power lateralization (i.e., alpha power at electrodes contralateral to the distractor cue relative to alpha power at electrodes ipsilateral to the distractor cue) was linked to behavioral performance. Power spectra for five subjects did not display obvious alpha-band peaks. These subjects were therefore excluded from the present analyses (n = 27). The mean peak alpha frequency (across subjects) was 10.95 Hz (Figure 9A).

**Figure 9.**
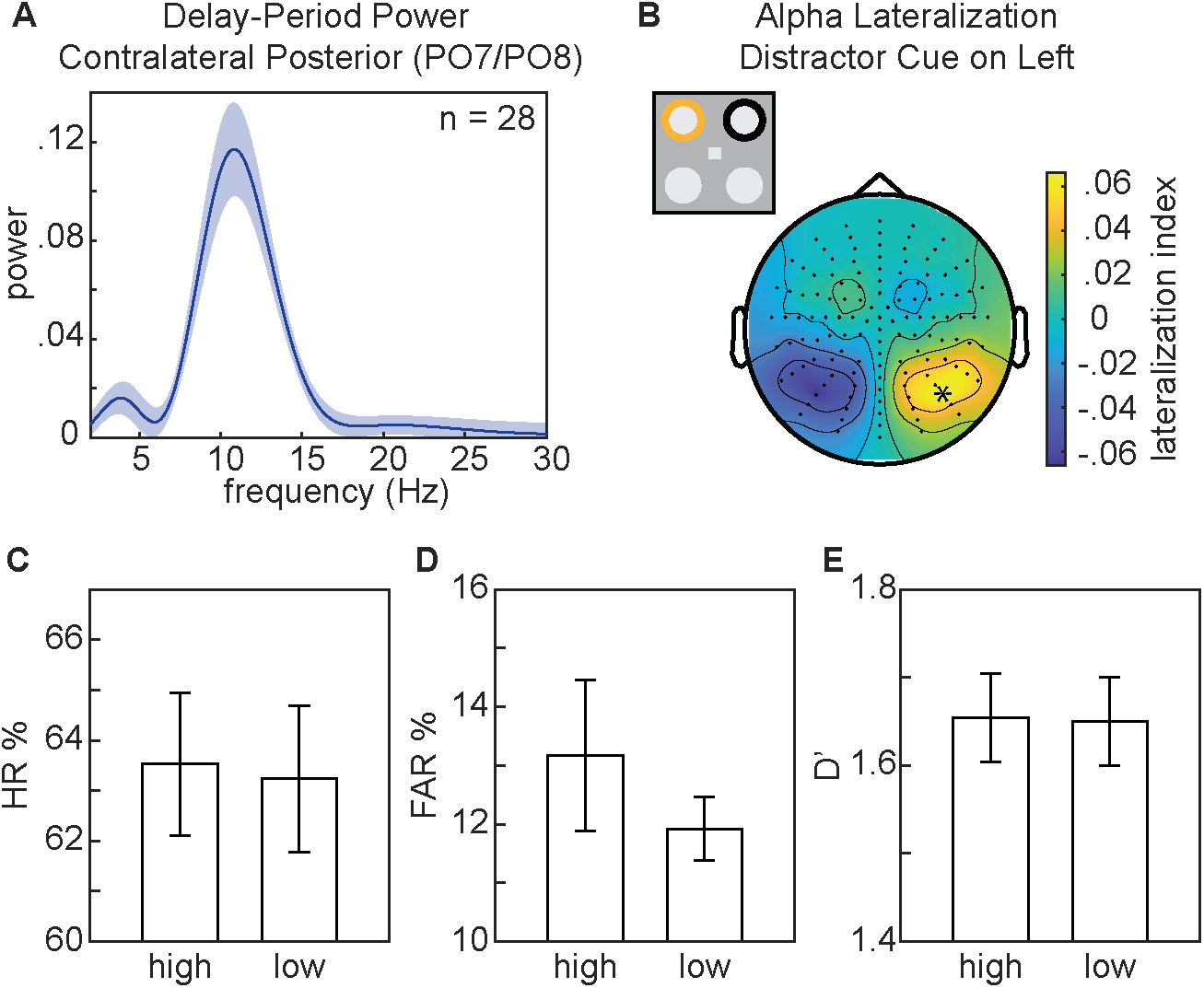
Posterior alpha power modulation does not affect task performance. ***A***, Delay-period (-500 to 0 msec relative to stimulus onset) power spectrum for frequencies from 3–30 Hz at occipital channels contralateral to the distractor cue. ***B***, Topography of alpha power lateralization relative to distractor cues presented on the left side. Channels were flipped across the midline for trials with the distractor cue on the right side. The asterisk indicates the channel used for binning trials by high and low alpha power lateralization. ***C***, Mean HR for trials binned by high and low alpha power lateralization using a median split. ***D***, Same as ***C*** but for mean FAR. ***E***, Same as ***C*** but for mean D’. All error bars indicate ±1 SEM.

The topography for alpha-power lateralization confirmed greater power over occipital electrodes that were contralateral to the distractor cue (Figure 9B), consistent with previous findings (Zhao et al., 2023); however, alpha-power lateralization at the occipital electrodes with the greatest alpha lateralization (A10/B7) was not associated with behavioral performance on trials with distractors [HR: *t*(26) = 0.164, *P* = .871, *d* = .032; FAR: *t*(26) = 0.892, *P* = .38, *d* = .135; D’: *t*(26) = 0.056, *P* = .956, *d* = .047] (Figure 9C-E). These results indicate that alpha-power lateralization in the present experiment—unlike alpha phase (see the previous analysis)—was not associated with suppression of distractor processing.

## Discussion

The present findings confirm that rhythmically occurring periods of worse target detection at a to-be-attended location are also associated with greater susceptibility to distractors (i.e., as indicated by higher false alarms), despite the location of those distractors being spatially predictable. These findings are consistent with the Rhythmic Theory of Attention (Fiebelkorn and Kastner, 2019), which proposes theta-rhythmic temporal windows associated with a greater likelihood of attentional shifts (i.e., theta-rhythmically occurring ‘shifting states’). Behavioral performance (i.e., HR and D’) on trials with and without distractors fluctuated as a function of pre-stimulus theta phase at central electrodes, peaking at ∼7 Hz. While theta phase was associated with fluctuations in distractor susceptibility, affecting FAR only on trials with a distractor, it was not associated with changes in distractor processing (i.e., it did not modulate the amplitude of distractor-evoked responses). In addition to theta-dependent effects, behavioral performance on trials with distractors further fluctuated with pre-stimulus alpha phase at both frontocentral and occipital electrodes, peaking at ∼9–10 Hz. Alpha-band effects (i) emerged exclusively on trials with distractors, (ii) were stronger at occipital electrodes that were contralateral to distractor locations, and (iii) modulated the amplitude of distractor-evoked responses. This pattern of results is consistent with an alpha-mediated gating of distractor processing (Jensen and Mazaheri, 2010). The present findings thus confirm a key prediction of the Rhythmic Theory of Attention—rhythmic susceptibility to distracting information—and provide evidence for distinct, frequency-specific attentional mechanisms that phasically modulate the influence of distractors on task performance.

A growing body of literature has demonstrated that spatial attention involves theta-rhythmic processes (for reviews see Fiebelkorn and Kastner, 2019; Benedetto et al., 2020; Fries, 2023; Re et al., 2025), revealing both theta-rhythmic modulation of neural responses (Busch and VanRullen, 2010; Landau et al., 2015) and theta-rhythmic modulation of behavioral performance (VanRullen et al., 2007; Landau and Fries, 2012; Fiebelkorn et al., 2013; Song et al., 2014; Dugué et al., 2015; Gallina et al., 2024). That is, there is a ‘good’ theta phase associated with better visual-target detection at the to-be-attended location and a ‘bad’ theta phase associated with worse visual-target detection at the to-be-attended location. Research in non-human primates has specifically linked these theta-rhythmic fluctuations in neural responses and behavioral performance to cortical and subcortical regions within the large-scale network that directs spatial attention, including frontal and parietal cortices (Fiebelkorn et al., 2018; Fiebelkorn et al., 2019; Gaillard et al., 2020; Spyropoulos et al., 2018). The Rhythmic Theory of Attention (Fiebelkorn & Kastner, 2019) proposes that theta-rhythmic fluctuations during attention-related sampling could promote critical cognitive flexibility, suggesting that temporal windows of relatively enhanced processing at the to-be-attended location (i.e., a ‘sampling state’) alternate with temporal windows associated with a higher likelihood of attentional shifts (i.e., a ‘shifting state’). This proposal is based on the specific cell types and neural dynamics that characterize alternating, theta-dependent attentional states (Fiebelkorn et al., 2018; Fiebelkorn and Kastner, 2019). Although such theta-rhythmically occurring windows of opportunity for shifting attention (i.e., proposed ‘shifting states’) might prevent us from becoming overly focused on any single location in the environment, these temporal windows might also make us more susceptible to salient information that can interfere with task performance (e.g., a distractor at a non-target location). The present results are consistent with this prediction, demonstrating that the theta phase associated with a lower HR (i.e., the proposed ‘shifting state’) was also associated with a greater distractor-related increase in FAR (Figure 7B). We are thus theta-rhythmically more susceptible to distracting information, meaning that windows of opportunity for shifting attentional resources can have behavioral disadvantages—at least in the presence of salient distractors. It should be further noted that theta-rhythmic fluctuations in distractor susceptibility occurred despite the location of the distractors being spatially predictable. That is, mechanisms of attention-related suppression—associated with a spatially informative distractor cue (Figures 3 and 4)— did not overcome the theta-rhythmic tendency to shift attentional resources (Benedetto et al., 2020; Dugue et al., 2016; Fiebelkorn and Kastner, 2019; Landau and Fries, 2012; Senoussi et al., 2019).

Previous research has shown that spatially predictable distractors can be actively suppressed to improve target detection elsewhere (Munneke et al., 2008; Chao, 2010; Chang et al., 2019; van Zoest et al., 2021; Redding and Fiebelkorn, 2024). The present experiment replicated such findings, demonstrating that spatially informative distractor cues promoted the suppression of distractors. That is, distractor cues were associated with both better visual-target detection (Figure 3) and lower-amplitude distractor-evoked responses (Figure 4). Previous research suggests that this process of active suppression—like attention-related sampling—might involve a neuro-oscillatory mechanism: specifically, neural activity in the alpha-frequency band (9–14 Hz) has been repeatedly linked to the suppression of sensory processing (Jensen and Mazaheri, 2010; Foxe and Snyder, 2011). For example, alpha power is typically higher in neural populations processing task-irrelevant locations and lower in neural populations processing task-relevant locations (Worden et al., 2000; Kelly et al., 2006; Thut, 2006; Yang et al., 2024). Further supporting this link between higher alpha power and the suppression of sensory processing, higher alpha power is associated with lower cortical excitability, as measured via spike rates and high-frequency band activity (Bonnefond and Jensen, 2015; Dougherty et al., 2015; Haegens et al., 2011; Spaak et al., 2012). Despite such frequently observed links between alpha-band activity and sensory suppression, results have been mixed regarding whether alpha-mediated mechanisms of spatial attention can be actively deployed to suppress spatially predictable distractors (Bonnefond and Jensen, 2025; Foster and Awh, 2019). Some studies have reported increased alpha power in anticipation of spatially predictable distractors, with associated effects on behavioral performance (Wostmann et al., 2019; Zhao et al., 2023) and distractor-evoked responses (van Zoest et al., 2021), while other studies have found no link between alpha power and the suppression of spatially predictable distractors (van Diepen and Mazaheri, 2017; Antonov et al., 2020; Redding and Fiebelkorn, 2024). The present results replicated an anticipatory lateralization of alpha power relative to the location of target and distractor cues, with higher power at electrodes contralateral to the distractor cue; however, the degree of this cue-related lateralization was not associated with behavioral performance (Figure 9). Here, we instead observed a significant relationship between all behavioral measures (i.e., HR, FAR, and D’) and the pre-stimulus phase of alpha-band activity (Figure 6). Previous research has similarly demonstrated a relationship between pre-stimulus alpha phase and behavioral performance (Mathewson et al., 2009; Dugué et al., 2011; Fakche et al., 2022; Zazio et al., 2022), but the present results specifically link these effects to distractor processing (also see Bonnefond and Jensen, 2012). That is, alpha-mediated fluctuations in behavioral performance only emerged on trials with distractors and were relatively stronger at occipital electrodes that were contralateral to distractors (Figures 6 and 8). These findings are therefore consistent with an alpha-mediated gating of distractor processing (Jensen and Mazaheri, 2010) that co-occurs with theta-rhythmic sampling and shifting (Fiebelkorn and Kastner, 2019).

Theta- and alpha-related effects in the present experiment were associated with different scalp topographies and different experimental conditions (i.e., alpha-related effects only emerged on trials with distractors). Figure 6 shows the topographies associated with phase-behavior relationships and indicates the electrodes that had significant effects, demonstrating a relationship between theta phase and behavioral performance during trials with and without a distractor (at central electrodes).

Topographic analyses revealed no between-condition differences for theta frequencies (6-7 Hz). In contrast, these analyses revealed significant differences between conditions for higher frequencies (8-10 Hz), reflecting alpha-band effects that were exclusive to trials with a distractor (at frontocentral and occipital electrodes). Furthermore, alpha phase—but not theta phase—was associated with fluctuations in distractor-evoked responses (Figures 7 and 8). The present results thus suggest that alpha-mediated mechanisms of spatial attention specifically influence the visual processing of distractors. Theta-mediated mechanisms of spatial attention, in comparison, might only indirectly influence distractor processing, by modulating the visual processing of targets (i.e., stimuli occurring at cued target locations). During the ‘shifting state’, weaker attention-related enhancement of sensory processing at the to-be-attended location could create a relative advantage for stimuli occurring elsewhere, thereby increasing the likelihood of attentional shifts toward salient distractors. Here, we were unable to test this hypothesis, because the low-contrast targets in the present experiment did not elicit a clear evoked response (i.e., we were unable to test whether the visual response to targets fluctuated as a function of oscillatory phase). The observed differences associated with theta- and alpha-mediated effects, however, are consistent with previous work that has proposed different functional roles for attention-related theta- and alpha-band activity, with theta-band activity perhaps being more closely associated with attention-related sampling and exploration and alpha-band activity perhaps being more closely associated with sensory processing and suppression (Dugué and VanRullen, 2017; Harris et al., 2018; Michel et al., 2022; Gallina et al., 2024; Galas et al., 2025).

The present findings (and those of others) provide evidence for theta- and alpha-mediated mechanisms of spatial attention, but it remains largely unknown whether and how these mechanisms interact. Several previous studies have described theta-rhythmic modulations of alpha power (Song et al., 2014; Fiebelkorn et al., 2018, 2019; Plöchl et al., 2022), with such interactions possibly being mediated through feedback projections from higher-order regions to visual cortex (Capotosto et al., 2009; van Kerkoerle et al., 2014; Marshall et al., 2015b, 2015a; Helfrich et al., 2017; Popov et al., 2017). Helfrich and colleagues (2017), for example, demonstrated that the phase of low-frequency oscillations in prefrontal cortex (∼2–4 Hz) modulated the effect of posterior alpha-band activity on behavioral performance. Here, we observed theta- and alpha-related effects with close proximity in the frequency domain (i.e., these effects occurred at similar peak frequencies), making it more difficult to conceive of a nested relationship, whereby theta phase modulates alpha power (Lakatos et al., 2005; Canolty and Knight, 2010). It might be possible, however, that theta-band activity in higher-order regions (e.g., frontal cortex) influences alpha phase in visual cortex (Dugué and VanRullen, 2017)—rather than alpha power— optimally aligning theta- and alpha-mediated mechanisms of spatial attention. Future research in non-human primates, which can measure interactions between different neural populations, will need to determine whether there are interactions between theta- and alpha-mediated effects on distractor processing.

The present results confirm that theta-rhythmic, attention-related sampling is associated with temporal windows of greater distractor susceptibility. That is, windows of opportunity for shifting attentional resources from the present focus of attention to another location increase the likelihood that a salient distractor will capture attentional resources. Rhythmic attention-related sampling—while perhaps preventing us from becoming overly focused on any single location—can also lead to behavioral disadvantages (i.e., increased distractor effects). These theta-rhythmically occurring fluctuations in distractor susceptibility co-occurred with alpha-mediated effects on distractor processing. Alpha-band activity contralateral to cued distractor locations phasically modulated both behavioral measures and distractor-evoked visual responses, consistent with an alpha-mediated gating of distractor processing. The present findings thus provide evidence for distinct theta- and alpha-mediated mechanisms of spatial attention, with both mechanisms phasically modulating the extent to which spatially predictable, high-contrast distractors interfere with the detection of spatially predictable, low-contrast visual targets.

## Acknowledgements

This work was supported by grants from the National Science Foundation (2120539), the National Institutes of Health (R01EY033726), and the Searle Scholars Program to I.C.F.

---

**Supplemental Figure 1.**
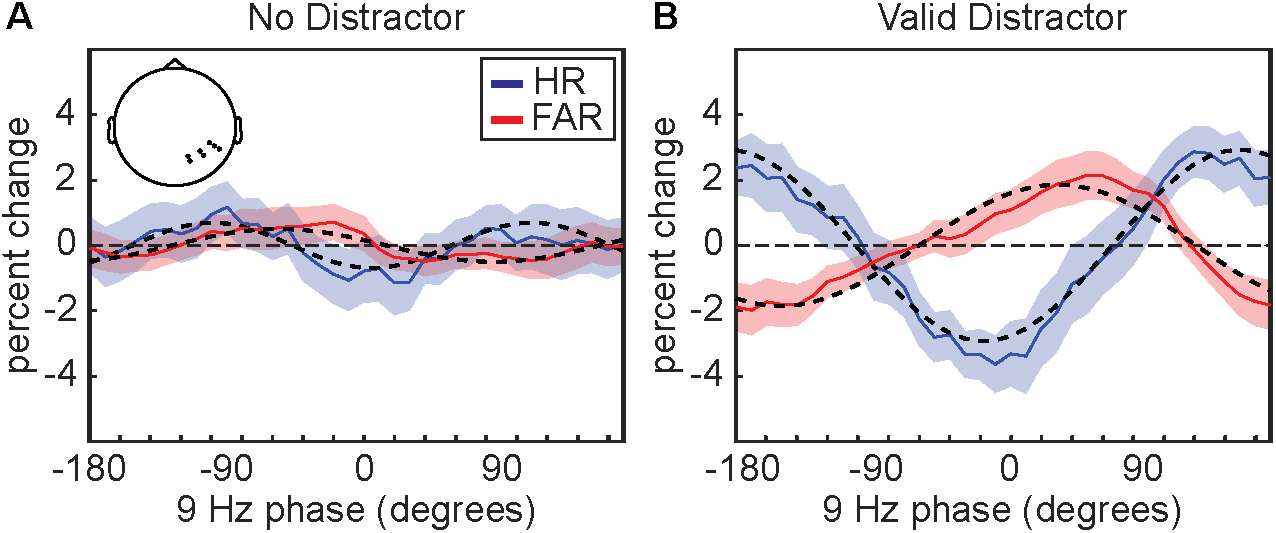
Alpha phase-behavior effects for posterior channels. ***A***, Phase-HR (solid blue line) and phase-FAR (solid red line) relationships for 9-Hz phase using trials with no distractor. These effects were averaged across posterior channels that were contralateral to the distractor cue and significant for trials with a valid distractor (depicted in inset). ***B***, Same as ***A***, but for trials with a valid distractor. All error bars indicate ±1 SEM.

